# Bridging Earth and Space: A Flexible and Resilient Federated Learning Framework Deployed on the International Space Station

**DOI:** 10.1101/2025.01.14.633017

**Authors:** James A. Casaletto, Patrick Foley, Mark Fernandez, Lauren M. Sanders, Ryan T. Scott, Shubha Ranjan, Shashi Jain, Nate Haynes, Marjan Boerma, Sylvain V. Costes, Graham Mackintosh

## Abstract

The public and commercial space industries are planning longer duration and more distant space missions, including the establishment of a habitable lunar base and crewed missions to Mars. To support Earth-independent scientific and medical operations, such missions can leverage artificial intelligence and machine learning models to assist with crew healthcare, spacecraft maintenance, and other critical tasks. However, transferring large volumes of data between Earth and space for model development consumes valuable bandwidth, is vulnerable to communication disruptions, and may compromise crew safety and data privacy. Federated learning enables model training while keeping data *in situ* and only transferring model parameters. In this work, we present a flexible, resilient federated learning framework that provides the secure transmission of model updates between Earth and the International Space Station. On March 15, 2024, this framework pioneered the deployment of federated learning in a spaceflight setting, training classifier models between Earth and the ISS using both real biomedical research data and synthetically generated data.

## Introduction

The National Aeronautics and Space Administration (NASA) and other space agencies have developed and deployed artificial intelligence and machine learning (AI/ML) for decades. These tools were used for purposes such as planning and scheduling space operations^1–3^, analyzing satellite images^4^, creating climate models^5^, detecting exoplanets^6^, identifying lunar features^7^, diagnosing anomalies^8–10^, and landing effectively on the surface of Mars in 2020 with the Perseverance Rover^11,12^. AI/ML models offer a powerful approach to support the needs of NASA and commercial spaceflight in its upcoming deep space exploration missions, such as Artemis, other portions of the Moon-to-Mars initiative^13,14^, and SpaceX’s StarShip missions. It is essential for deep space missions to develop capabilities for *in situ* analysis and greater crew autonomy, as these missions will lack immediate access to the continuous support of mission control staff, scientists, and medical personnel allocated to low Earth orbit (LEO) missions. To stay current, these models would need to be routinely updated with new data from Earth, ensuring they reflect the most advanced knowledge and insights available. However, large-scale data movement poses significant challenges, as ground-to-space data transmission is less reliable and slower than terrestrial networks. The time and cost associated with transferring large volumes of data can be prohibitive, and the high communication latency in deep space missions further complicates timely updates^15^. The small number of astronauts makes true anonymization challenging^16^, and more generally, open scientific research into the effect of spaceflight on the health of public and commercial astronauts is restricted by governmental policies^17–19^.

Federated learning is one solution to enable the training of machine learning models using Earth-based and space-based data without having to transfer data at all. This approach has seen advancements terrestrially across various clinical research approaches and applications^20–23^. Federated learning is an approach to building machine learning models when the data used to train, test, and validate them are stored on participating compute nodes in separate locations^24–26^. Each participating node is sent a copy of a machine learning model to train on their local data. Once the model has been trained locally over some number of iterations, the nodes send only their updated version of the model parameters to a central coordinating server without sharing their local data. The server uses some form of statistical aggregation (e.g., weighted or unweighted mean, or median) on the contributions from all the nodes and updates the global model. The updated parameters of the global model are shared again with all the nodes, and the process repeats until the global model converges. Since its inception, federated learning has emerged as a promising approach in machine learning when the training data are distributed and too large to consolidate or private and too sensitive to release for external use^27–30^.

Here we present the first machine learning models trained in a federated manner using data located on Earth and data located separately on HPE’s (Hewlett Packard Enterprise) Spaceborne Computer-2 (SBC-2) aboard the International Space Station (ISS)^31^. We introduce in this work a novel software framework termed FLUID (Federated Learning Using In-space Data) which leverages Intel’s OpenFL federated learning implementation^32^. A significant component of this work was dedicated to building the infrastructure required to enable the secure and efficient exchange of model parameters between Earth and the ISS, allowing updates to the model on both sides without transferring raw data. In the current study, we train the Causal Relation and Inference Search Platform (CRISP) machine learning ensemble^33,34^, but FLUID is model-agnostic and can be leveraged to train any type of machine learning model in a federated manner.

## Methods and Data

### FLUID architecture

Most distributed federation platforms, including OpenFL, operate within a Transmission Control Protocol/Internet Protocol (TCP/IP) network environment^35^. Client collaborator applications make TCP or User Datagram Protocol (UDP) connections to a central aggregation server and provide their model updates immediately. At the simplest level, this typical deployment is depicted in Figure 1.

**Figure 1:**
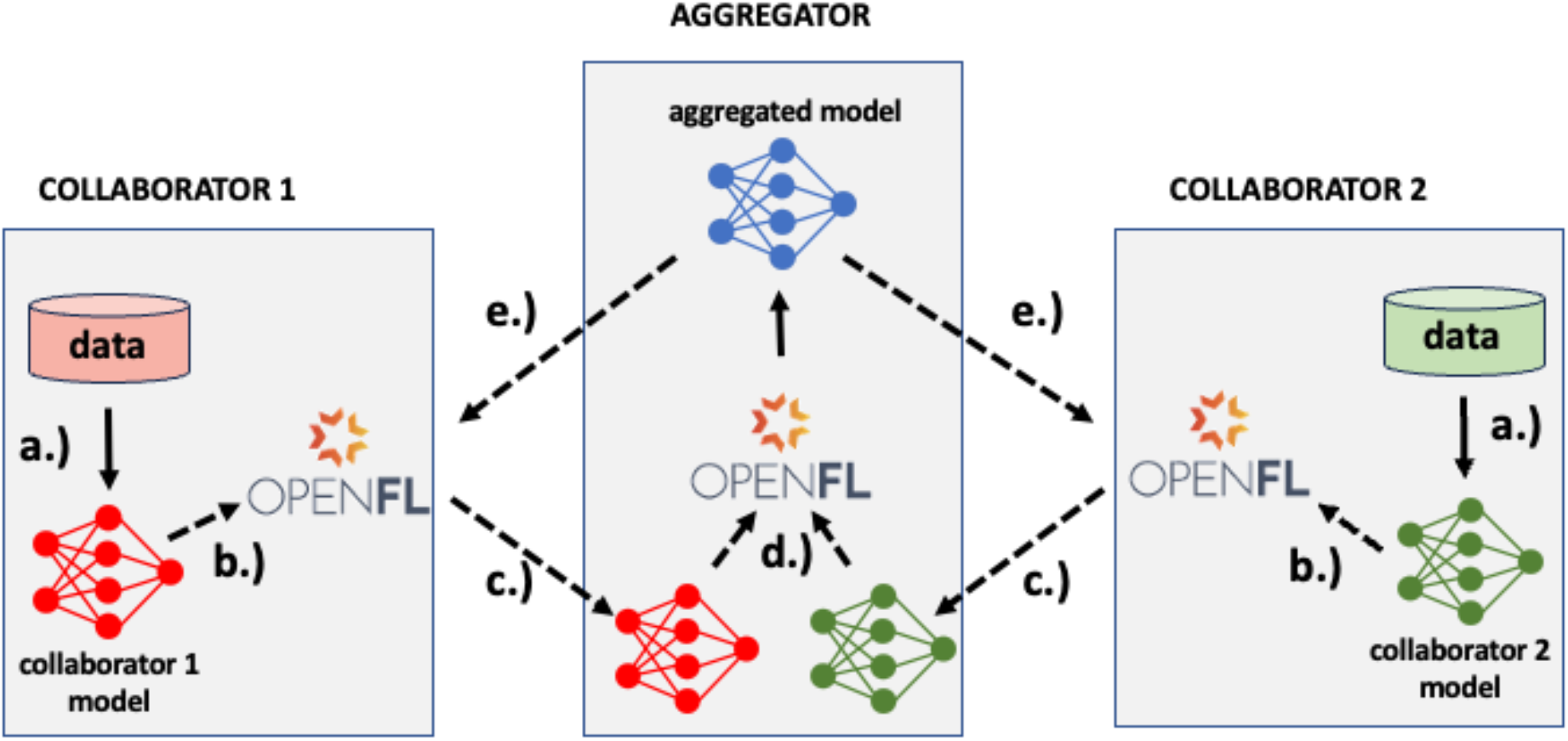
Typical OpenFL architecture. a.) OpenFL collaborators train a local model based on their local data. b.) At periodic intervals, the training algorithm stops and sends the current version of the model parameters to the OpenFL library. c.) OpenFL collaborators send the model parameters to the aggregator over a TCP/IP network. d.) OpenFL aggregator merges (averages) the model parameters from all the collaborators to create a single global model. e.) OpenFL aggregator sends the global model to all collaborators to replace their local model with the global one.

In a typical OpenFL architecture, communication is exchanged over a TCP/IP network connection. However, TCP/IP connections are not persistent or stable between Earth and space, so modifications were made to the OpenFL architecture and to the unique communication protocols between SBC-2 on the ISS and its ground control Earth terminal. These modifications abstracted and encapsulated the complexities of communications with the ISS and SBC-2. OpenFL uses the Google Remote Procedure Call (gRPC) framework^36^ to enable distributed communication between collaborators (clients) and the aggregator (server). Internally, gRPC uses protocol buffers (protobuf) as a data format to exchange between clients and servers. The protobuf data format is an extensible, language- and platform-neutral method for serializing structured data, making it suitable for communications protocols, data storage, and various other applications^37^. Because protobuf can also be used as a file format, there is no need to translate the data or to use other protocols. The protobuf data can be read from files and written to the network. Similarly, protobuf data can be read from the network and written back to files. This is precisely what our modifications entail: the addition of shim nodes, so-called because they each act as half of a 2-way relay between Earth and space, which read data from and write data to network and files. There is a node called an aggregator shim which runs on Earth, reads files from a file system in protobuf format, and sends the protobuf messages over a TCP/IP connection to the Earth aggregator using the normal OpenFL protocol, gRPC. The ISS collaborator sends messages using the normal OpenFL protocol over a TCP/IP connection to the ISS collaborator shim (the other half of the 2-way relay between Earth and the ISS) which, in turn, writes these messages to files in the file system. Consequently, the rest of the OpenFL platform, and the federated learning application being trained on it, could be developed, deployed and executed in a completely standard manner, as if the SBC-2 were terrestrially networked instead of being in orbit. Figure 2 depicts the entire data flow in the FLUID architecture between the nodes running in Amazon Web Services (AWS) on Earth and those running in space on the Hewlett Packard Enterprise (HPE) SBC-2.

**Figure 2.**
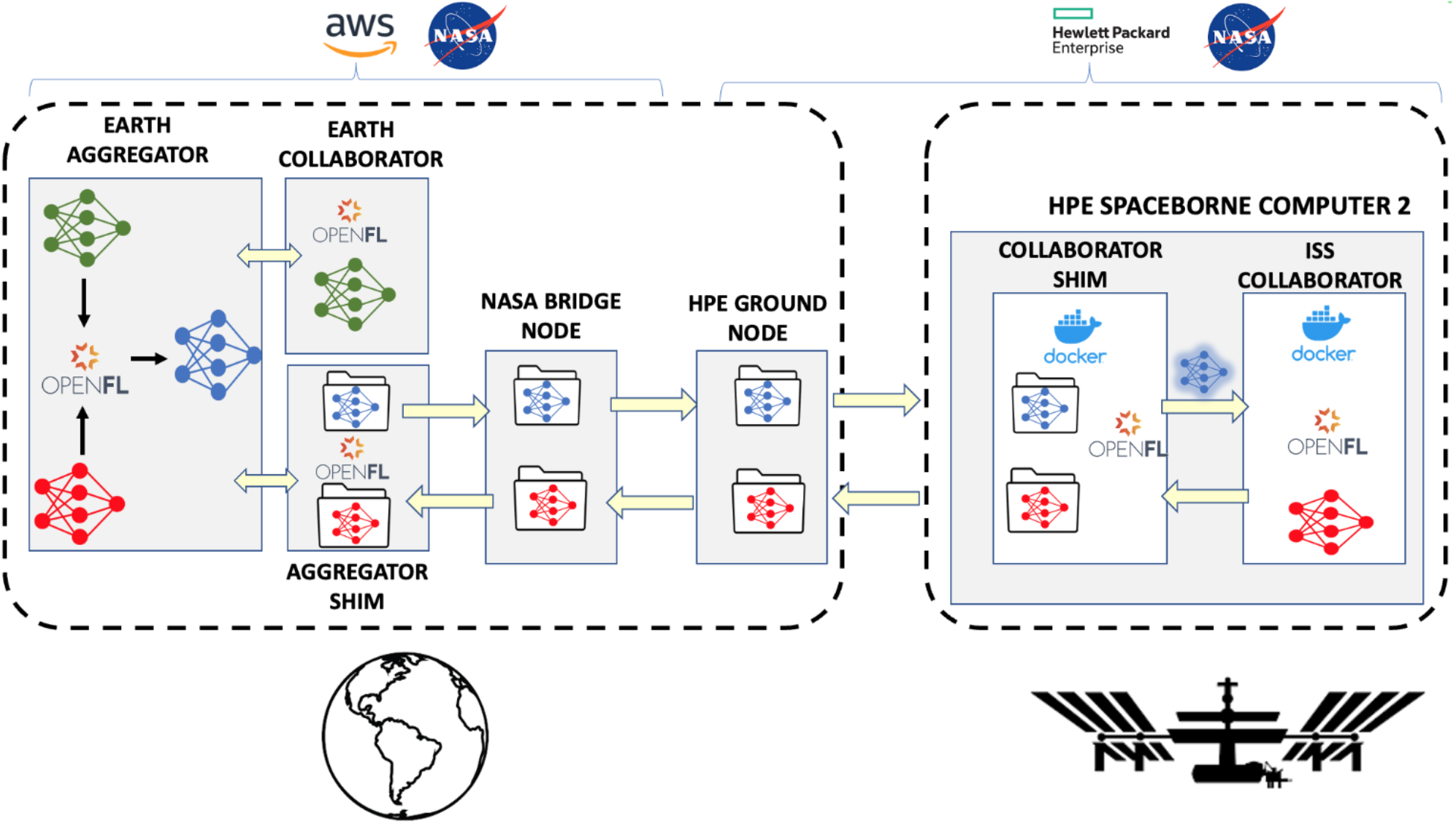
Data flow between collaborators and aggregator in ground-to-space federated model training. After each round of training, the Earth and ISS collaborators build a model (shown in green and red, respectively) based on their local data. The Earth collaborator sends its model directly to the Earth aggregator over TCP/IP, while the ISS collaborator sends its model to the collaborator shim which, in turn, writes the model to a file. This file gets transmitted down to the HPE ground node, transferred to the NASA bridge node, and sent to the aggregator shim which reads the model from the file and sends it to the Earth aggregator over a standard TCP/IP connection. Finally, the Earth aggregator combines these two collaborator models to build a global model (shown in blue). The global model is then sent back to the Earth collaborator (via TCP/IP) and ISS collaborator (via the shim nodes) to use in the next round of training.

The main data flow complexities that this FLUID architecture addressed were two-fold. First, frequent and lengthy loss-of-signal (LOS) events occur between the ISS and Earth communication stations (i.e., complete communication outages for several minutes). Unpredictable losses of signal can occur as well, and the FLUID platform must tolerate these. Second, the communication protocol between the SBC-2 and the ground control terminal is limited to batch synchronization of files which emulate a live TCP/IP network connection between the Earth-based aggregator and the collaborator in orbit.

The communication link between Earth and the ISS undergoes predictable signal loss due to the alignment of terrestrial satellite towers and the ISS’s orbital orientation. They also occur during maintenance windows and hardware or software failures. When the communication link between Earth and the ISS is unavailable, the FLUID shim nodes simply buffer the files that are being generated on Earth and on the ISS until the communication link is established again. During LOS events, the collaborators synchronously and indefinitely block program execution until they get a response back from the aggregator which contains the newly updated model. In this way, collaborators can tolerate arbitrarily long communication outages and then continue exactly where execution left off prior to the LOS.

Experiments which run on the SBC-2 must execute inside Docker containers. To manage running the federation, we developed shell scripts that launch Docker containers in AWS and on the ISS. These scripts read the OpenFL experiment configuration from files, mount the necessary data volumes that contain the training and test data, and start the OpenFL federated framework on the collaborators, aggregator, and shim nodes. Additionally, we developed scripts to connect the data flow from the AWS bridge node to the HPE ground node (also known as the test and development system; TDS) where data is exchanged with the ISS. The bridge node and HPE ground node scripts use the rsync command to synchronize files generated on Earth and in space. Because the HPE ground node is protected by a firewall, the bridge node IP address was fixed and added to the ingress firewall rules of the TDS.

### CRISP v2.0 machine learning ensemble

We utilized CRISP v2.0, the second iteration of the Causal Research and Inference Search Platform which was developed for the 2021 Frontier Development Lab Astronaut Health challenge. CRISP v1.0 comprises an ensemble of causal inference algorithms that identify a subset of explanatory variables most likely to have a causal impact on a target binary variable. CRISP v2.0 uses OpenFL to bring federated learning to the ensemble, enabling researchers to conduct causal inference across decentralized data sources without necessitating the exchange of data. CRISP v2.0 can run in Docker containers, abstracting the operating environment from the underlying physical host. This software is a key contribution to the existing work on edge computing, cloud computing, and AI/ML applications for space biosciences^38,39^.

### HPE Spaceborne Computer

HPE’s SBC is a collaboration between HPE and the International Space Station National Lab, with support and involvement from NASA. The mission of SBC is to demonstrate the feasibility and value of using modern, unmodified hardware and software for high performance computing, edge processing, and AI/ML scientific experiments off planet. HPE’s third iteration of Spaceborne Computer launched January 30, 2024.

This configuration of the award-winning HPE Spaceborne Computer, based on HPE Edgeline and ProLiant servers, has been updated with over 130 TB of flash-based storage from KIOXIA, the most storage to ever travel to the ISS on a single mission. This includes four KIOXIA 960 GB RM Series Value SAS, eight 1,024 GB XG Series NVMe and four 30.72 TB PM6 Enterprise SAS SSDs. The additional flash memory storage makes it possible to run new types of applications and conduct scientific research using larger data sets. Specifically for the FLUID scientific experiment, the HPE Edgeline EL4000 component of the Spaceborne Computer was used. The HPE Edgeline 4000 runs the standard, unmodified Red Hat 7.8 operating system using a single socket x86 system and a single Nvidia T4 GPU. The SBC EL4000 system is connected via 1GbE to the ISS Local Area Network for communications back to Earth and into HPE’s SBC secure lab. Also within HPE’s SBC Lab, and within HPE’s firewall, are identical systems for certifications for flight. These are analogously called Earthborne Computer (EBC). Additionally, within the SBC Lab is a Command and Control System which can safely and securely access the SBC systems on the ISS and identical systems outside the HPE firewall (the TDS). Partners are granted limited and restricted access to these TDS systems.

Once a science experiment has been successfully run on a TDS system, it is moved to the Earthborne systems for flight certifications. It is at this point that the network characteristics of ISS communications are simulated for flight certification using a secure air-gapped environment. Once successful, the updated experiment is installed on SBC aboard the ISS. For the FLUID experiment, the aggregators, collaborators, shims and other components as described above were first developed on AWS. Next the appropriate modules were ported to the TDS systems. Once successfully tested on the TDS systems, they were installed on EBC Systems. Finally, after further testing, certifications and NASA approval, FLUID was installed on SBC-2 aboard the ISS. These Docker containers are unaware of and unable to handle intermittent data communications. For these Docker containers to exchange data within this highly secure, multi-system environment, we developed scripts to monitor connectivity, cache data, preserve permissions, update ownerships and verify correctness in order to maintain a seamless data flow to and from the edge of the edge on the ISS and the Earth-based AWS. With these scripts, we have successfully encapsulated and abstracted ground-to-space communications in a safe and secure manner.

The Telescience Resource Kit (TReK) is a suite of software applications and libraries that can be used to monitor and control assets in space or on the ground^40^. The Huntsville Operations Support Center (HOSC) Payload Ethernet Gateway (HPEG) service is accessible through the TReK HPEG application. The HPEG service provides access to payloads onboard the ISS using standard network protocols and services. Services supported include ping over Internet Control Messaging Protocol (ICMP) and Secure Shell (SSH) over TCP port 22. The SSH service enables sending shell commands to the SBC-2 for running experiments such as this, and pings sent at regular intervals help determine network connectivity between Earth and the ISS. Figure 3 shows a screen capture of the HPEG tool displaying LOS events between Earth and the ISS.

**Figure 3.**
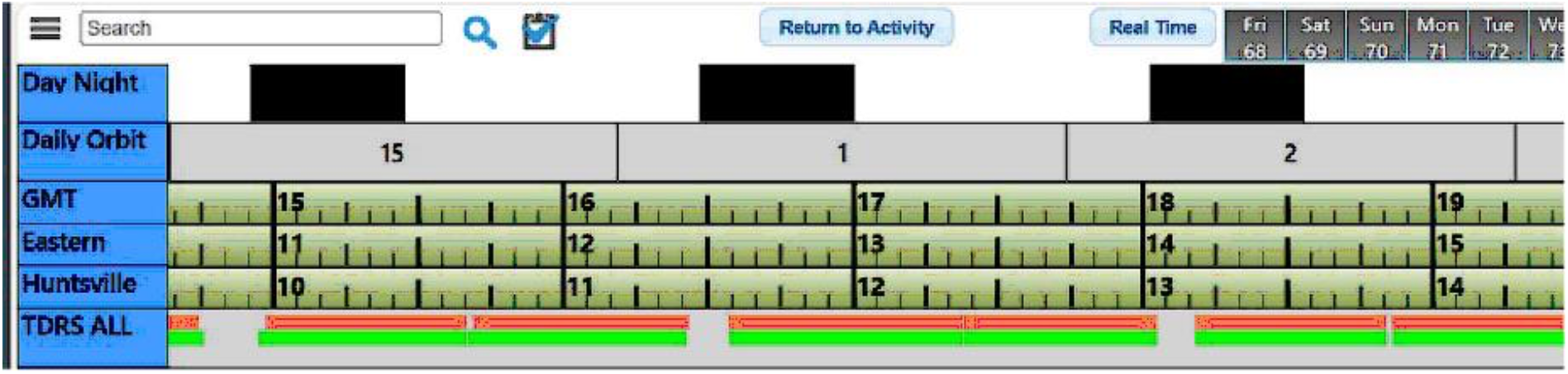
Loss-of-signal windows between HOSC and the ISS. Known LOS windows displayed in the HPEG user interface. The ISS communicates with Earth through a robust network of ground stations and geostationary satellites called the Tracking and Data Relay System (TDRS). Due to predictable line of sight constraints, there are brief gaps in coverage, as shown in the “TDRS ALL” row of this image.

### Synthetic data

We conducted two use-case federation exercises on the ISS: one with synthetic data and one with biomedical data. The synthetic data were generated using utilities in the CRISP software repository. These utilities leverage structural equation modeling (SEM) to embed a known causal structure in a tabular dataset whereby specific feature values were configured to be the features “causal” of a target variable^41^. SEM is a collection of statistical techniques that allows a set of relationships between one or more independent variables, either continuous or discrete, and one or more dependent variables, also either continuous or discrete, to be analyzed. Using SEM, we generated a synthetic dataset with a target variable, 30 features with a mathematically causal relationship for predicting the target, and 320 features that are not at all related to the target. We generated random feature values in the causal columns by drawing from a standard normal distribution (mean 0 and variance 1). The target value was first generated using a function of the causal feature values, squashed to the unit interval [0, 1] using a sigmoid function, and then binarized to either 0 (if < 0.5) or 1 (if >= 0.5). There were a total of 1000 samples in the synthetic dataset which were randomly split 50/50 between the Earth-based collaborator and the ISS-based collaborator. At each site, the 500 data points were split into 70% for training and 30% for testing.

### Cardiac data

In addition to using synthetic data, we also ran an experiment using real biomedical data from the NASA Open Science Data Repository (OSDR)^42,43^. The OSD-435 dataset^44^ contains data on function and structure of the heart and aorta in mice exposed to various types and doses of ionizing radiation. Cardiac function and blood velocity were measured with ultrasonography at 3, 5, 7, and 9 months after irradiation, the results of which were published by Seawright et al^46^. These non-invasive echocardiography techniques could be deployed in the future on a deep space mission to collect similar measurements from astronauts and model organisms. The OSD-435 dataset serves as a simulation of a situation in which cardiovascular biomedical data are collected both in space and on Earth and used to train a predictive machine learning model in a biomedical scenario. The data were filtered to remove results from low-dose radiation (0.05-0.25 Gy) to reduce noise and filtered to remove samples without a number in any field. The dataset features used as predictors in the model include radiation type, time (in months) post-irradiation, heart rate, peak velocity, velocity-time integral, and mean velocity. The binary targets in the CRISP classifier were derived from the dose values with 0 Gy in class 0 and >=0.5 Gy in class 1. The dataset was split 70/30 train/test, and 50/50 between collaborator 0 and collaborator 1.

## Results

We performed two experiments using the FLUID federated learning platform in order to demonstrate the ability of federated learning to train machine learning models using data on Earth and in space. For the first experiment, we used synthetically generated data in which the features with a causal relationship to a target variable were mathematically generated, providing known ground truth (see Methods). For the second experiment, we used data on cardiac function in mice exposed to simulated space radiation from the OSD-435 dataset.

### Federated model trained with synthetic data

The CRISP v2.0 ensemble was trained to identify causal features of the target variable in the synthetic dataset for which causality was mathematically established via the SEM approach (see Methods). The dataset was split 50-50 between collaborator 0 (on Earth) and collaborator 1 (in space), and the CRISP v2.0 model was trained as a binary classifier (causal vs. non-causal). As a binary classifier, the null model performance is 50% accuracy. Figure 4 shows the trajectory of the CRISP binary classifier ensemble accuracies over the course of the training rounds for both the federated and centralized model training.

**Figure 4.**
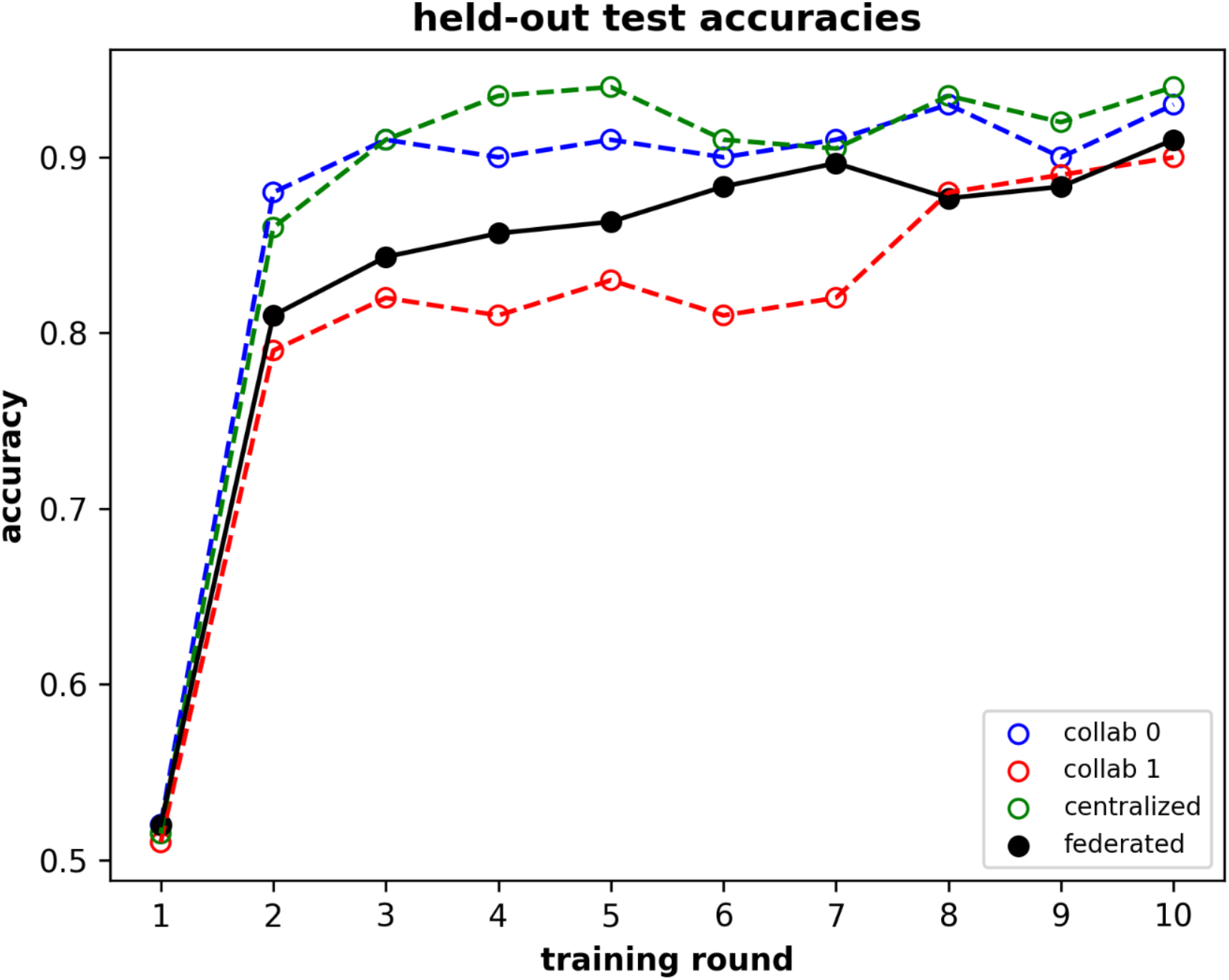
Test accuracies for the synthetic training dataset. Federated test accuracies are shown in black, and centralized test accuracies are shown in blue (model using data only from collaborator 0), red (model using data only from collaborator 1), and green (model using data concatenated from both collaborators). After 10 rounds of training with a single epoch each, the test accuracies of the models trained on this dataset converge within 5% of each other.

The federated learning and centralized learning models demonstrated similar accuracy. In general, the performance of a federated learning deployment compared to centralized learning depends in part on the distribution of the data amongst the collaborators. Ideally, the data on all the collaborator nodes is independent and identically distributed (IID), but this is often not the case. Such non-IID datasets can result from biases and discrepancies in data generating methods across various sources. This divergence from the expected global distribution can decrease the overall performance of the federated model.

Another factor that contributes to federated model performance is the amount of data that is available on each of the collaborators. In our ISS synthetic data experiment shown in Figure 4, we split the data 50-50 between the two collaborators. After that experiment, we conducted additional experiments on Earth to observe the effect of data distributions on model performance. Figure 5 shows the federated model performance for four different synthetic data distributions between the collaborators.

**Figure 5.**
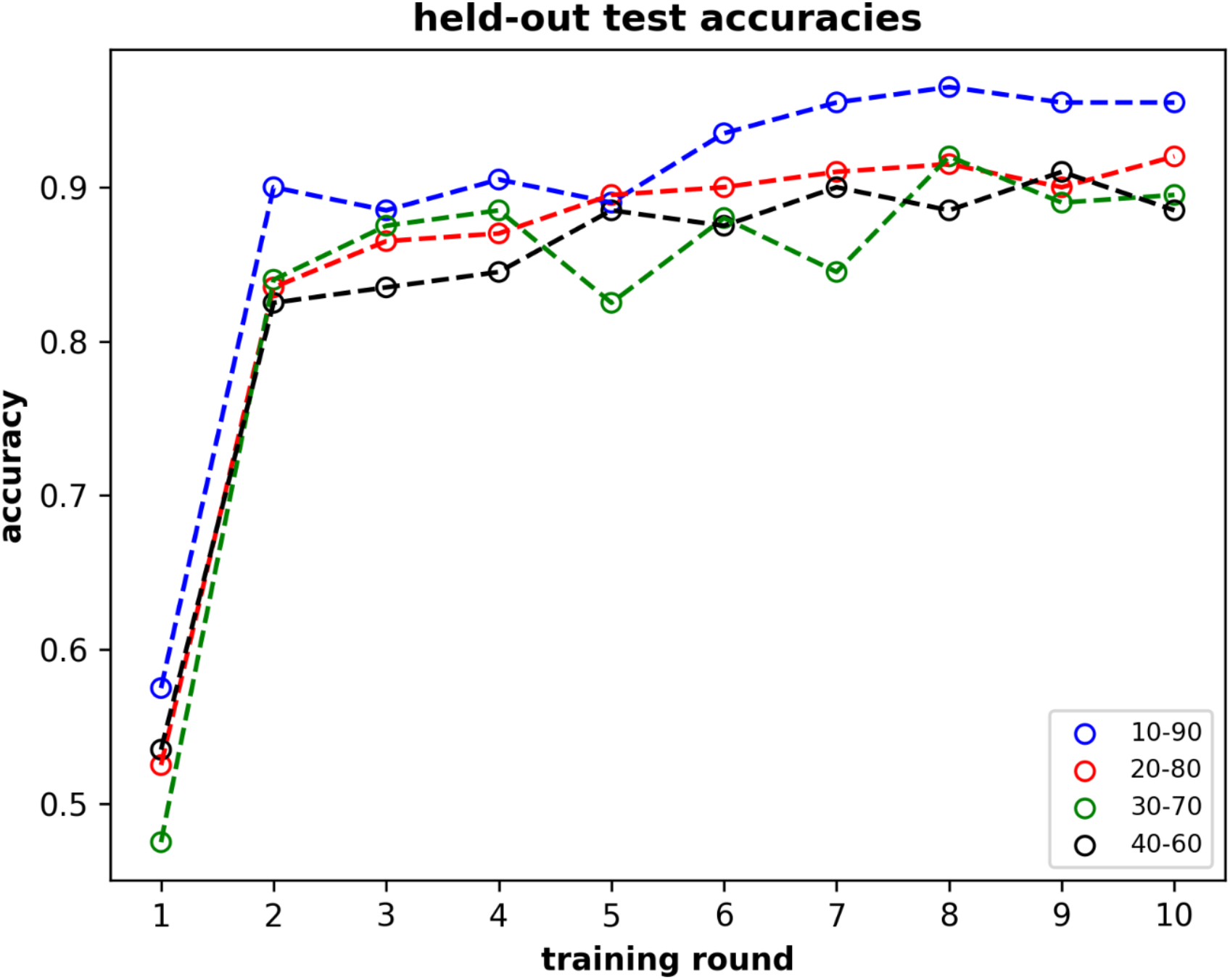
Federated model test accuracies for different distributions of the synthetic training dataset. The 10-90 split of data yields the best overall performance and the 40-60 split of the data yields the worst, suggesting that the averaged model parameters weighted by data size can overcome data size discrepancies between collaborators.

### Federated model trained with space biology data

To demonstrate the potential of the FLUID platform for training and predicting on biomedical measurements from astronauts during deep space travel, we simulated a situation in which cardiac data were collected on the ISS and used to train a model enhanced with relevant cardiac data from a model of simulated space radiation exposure on Earth. We split the dataset 50-50 between collaborator 0 (on Earth) and collaborator 1 (in space) and trained a model to identify which features were most causally related to radiation exposure. Interestingly, blood velocity was most closely linked to radiation dose, identifying a potential relationship between radiation exposure and cardiac function. Figure 6 shows the trajectories of accuracies over time for models built centrally and using FLUID.

**Figure 6.**
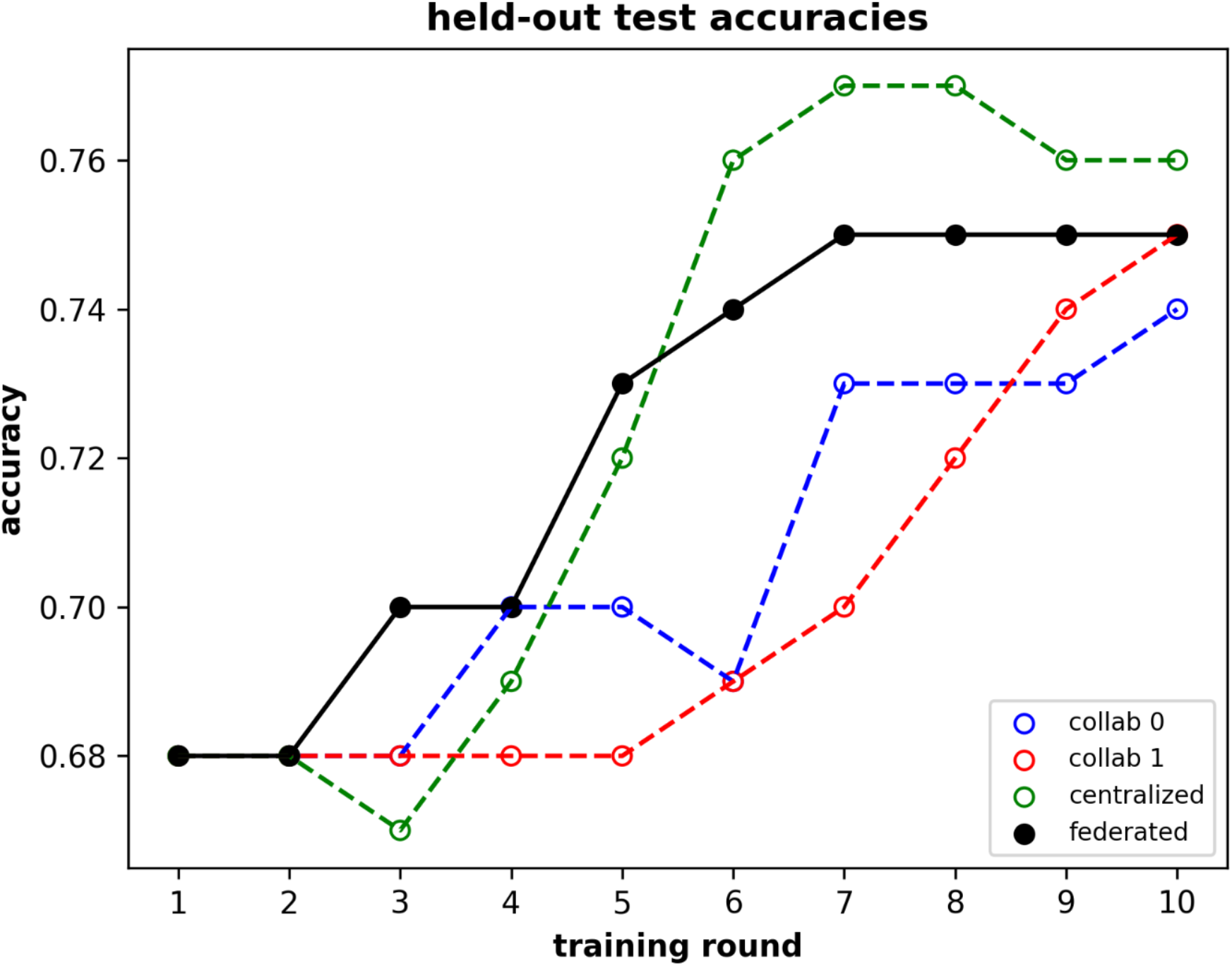
Test accuracies for the space biology training dataset. The blue plot depicts the progression of accuracy for a centralized model trained only on data assigned to collaborator 0, and the red is the progression of accuracy for a centralized model trained only on data assigned to collaborator 1. The green plot depicts accuracy for a centralized model trained on all the data located in one location, and the black depicts accuracy for a federated model trained with 2 collaborators.

Because the data came from the same experiment and each sample’s measurement is independent of all other sample measurements, the data are IID. Therefore, the discrepancies in model accuracies are due at least in part to the random nature of how the data were split. If these data were separately generated on Earth and in space, differences in the echocardiography instruments could introduce biases between multiple collaborators. We could then leverage the techniques discussed in the Methods section to harmonize the non-IID data. This simulation demonstrates the ability to train a model using astronaut or model organism cardiac measurements on a deep space mission, together with cardiac data from Earth-based animal models or human subjects, to increase predictive power.

### FLUID tolerates loss of signal events

The FLUID architecture is designed to tolerate LOS events that last arbitrarily long. Figures 7a and 7b depict the Earth-ISS communication status in Greenwich Mean Time (GMT) from HPEG just after the LOS and the acquisition of signal (AOS), respectively. Figure 7c shows the collaborator shim node program output during and just after an LOS event which was experienced when running this experiment.

**Figure 7.**
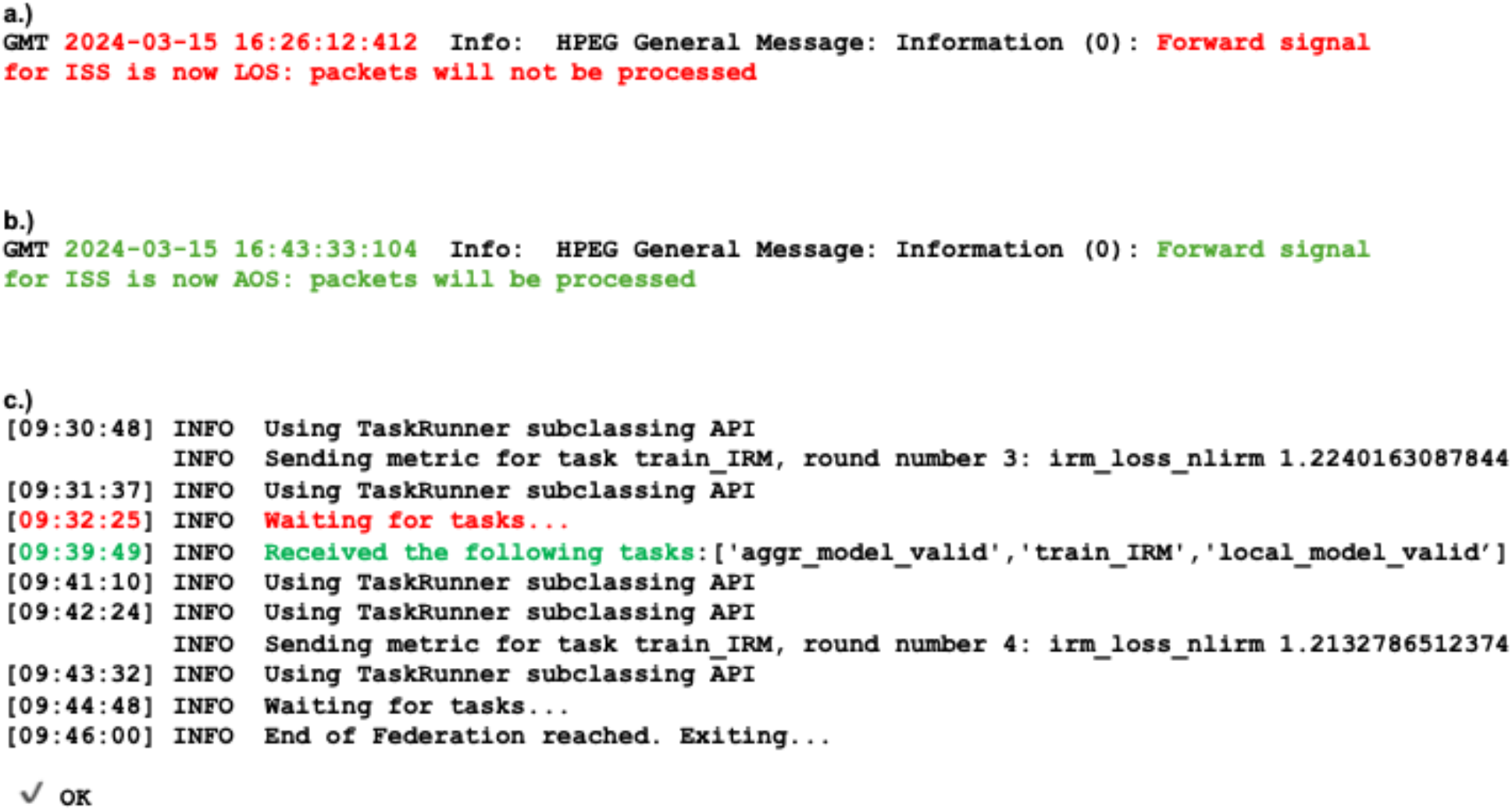
Federated training persists through LOS. a.) Log message from HPEG denoting the start of the LOS (in GMT). b.) Log message from HPEG denoting the end of the LOS (in GMT). c.) Output from the collaborator shim node running on the SBC-2 shows a pause (“Waiting for tasks…”) during the LOS from 9:32:25 to 9:39:49 (Pacific Time).

## Discussion and Conclusion

In this study, we present the FLUID federated learning platform and report the first successful implementation of machine learning models trained in a federated manner using data separately held on Earth and aboard the ISS. The successful completion of this work has far-reaching implications for AI/ML utilization during deep space missions. The ability to train predictive models with federated data opens the door for leveraging Earth-based datasets to generate insights and predictions during deep space travel. For example, the FLUID platform could train a model to predict variations in astronaut heart rates by using billions of data points from wrist wearables on Earth and refining the model with heart rate data from astronauts during space flight or while living in a space habitat. Our results demonstrate that this software can train predictive models with accuracy comparable to centralized models without requiring the transfer of potentially sensitive or large datasets between Earth-based and space-based computers. Additionally, the system effectively recovered from extensive communication interruptions caused by multiple LOS events during model training. While the current study demonstrates the potential of federated learning for space exploration, the demonstration is limited to datasets artificially split between space and Earth. In a realistic federated learning scenario, an aggregator would be trained on data generated on a spacecraft, habitat, or cubesat, while one or more additional aggregators would be trained on large datasets residing on Earth. Only model weights would be shared with space-based systems, enabling the prediction of space-relevant phenomena after baseline training on Earth-based databases.

Federated learning would provide significant advantages for space travelers. These include enabling predictive model training that leverages the vast data resources on Earth without requiring the transfer of large datasets to spaceborne computers aboard spacecraft or habitats^47^. To enable federated learning on deep space missions, future work should prioritize improving federated learning capabilities to facilitate appropriate data sharing and preprocessing methods among collaborators. A critical area for future exploration is developing techniques to assess and mitigate the impact of non-IID data provided by collaborators. Non-IID datasets often arise from discrepancies in data collection methods across different sources, leading to deviations from the expected global distribution and reducing overall federated model performance. Promising methods for data harmonization across collaborators include standardization, normalization, batch effect correction, and domain adaptation^48–50^.

With the advent of large language models (LLMs), their integration with active learning architectures, and high-performance computing in space, it is now feasible to send predictive models to space pre-trained on high-resource Earth databases and update them using space-generated data. These models could operate in diverse environments, such as LEO, lunar missions, Mars transit, or the Martian surface. Such advancements would enable astronauts to access all relevant knowledge from Earth without requiring communication with Earth during high-latency situations or LOS events. For instance, these models could support a Precision Space Health system that autonomously monitors and evaluates biomedical data, predicts health risks, adapts to new information, and provides timely insights and decision support for deep space crew members and their chief medical officer^47^. The FLUID platform could refine a pre-trained biomedical question-answering LLM with space-relevant documents from NASA, other space agencies, and private companies. Astronauts could query this LLM during communication blackout periods to address urgent medical questions, serving as a clinical decision support tool^51^. Furthermore, FLUID could facilitate fundamental life sciences research in space. Automated “self-driving” laboratories could generate experimental data to study the effects of spaceflight in high-throughput, intervention-free settings^52^. This data could train predictive models while simultaneously leveraging extensive public life sciences databases on Earth.

A critical aspect of FLUID’s application is the need for longitudinal data updates to ensure models remain relevant over extended missions. Models could be updated at specific cadences, balancing the benefits of incorporating new insights with the challenges of transmitting and integrating data across vast distances. Data storage considerations are also essential, particularly for managing limited storage capacity on spacecraft. As models evolve, decisions must be made regarding the “aging” or “retirement” of data that is no longer relevant. Addressing these issues is key to avoiding *in situ* data overload, which could hinder computational efficiency and model performance^52^.

Approaches like FLUID enable space travelers to access the wealth of terrestrial biomedical and scientific data without the need to transfer it to spaceborne systems (Figure 8). Instead, federated learning ensures that only model weights are transmitted, dramatically reducing the burden on communication networks. This also supports the efficient use of storage and computational resources on spacecraft.

**Figure 8.**
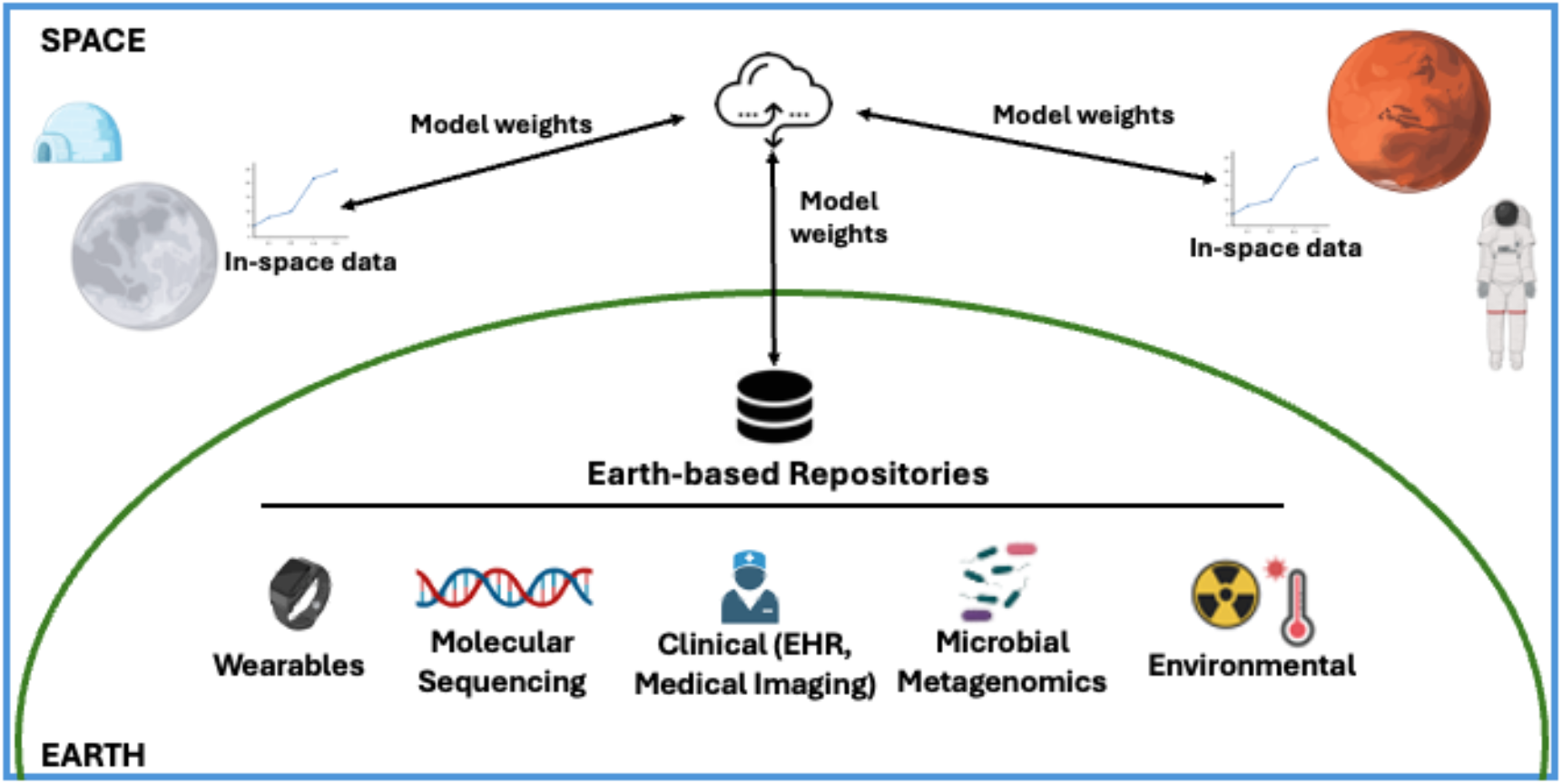
FLUID enables space travelers access to Earth-based repositories. Federated learning approaches like FLUID enable light-weight updates to spaceflight predictive models, leveraging the wealth of knowledge in Earth-based repositories while only transferring model weights instead of raw data.

FLUID could be leveraged for building models based on data generated across the solar system, from Earth and LEO to the moon, Mars, and beyond. Overall, FLUID represents a groundbreaking and novel approach, marking the beginning of a paradigm shift in enabling AI/ML model training and prediction in a federated manner during space missions. It bridges terrestrial and spaceborne data resources, significantly advancing edge computing, cloud computing, and AI/ML applications for deep space exploration, ultimately enabling humanity to become permanently multi-planetary.

## Author Contributions

GM, SJ, and PF designed the FLUID architecture and wrote some of the methods section. PF made all the software changes to OpenFL and wrote that part of the methods section. JC wrote all the scripts, built the containers, designed, ran, and analyzed the experiments, and wrote parts of the abstract, introduction, methods, and results sections. RS and LS identified the OSD-435 dataset for the experiments and wrote the data section and part of the abstract, introduction, and conclusion. SC wrote part of the introduction and conclusion. SR, NH, and GM made the AWS compute resources available. MF made the EBC and SBC-2 compute resources available, ran the experiments on the ISS, and wrote part of the methods section. MB ran the experiments which produced the OSD-435 dataset. All authors reviewed the manuscript and provided valuable feedback.

## Acknowledgements

This work was funded by the NASA Biological and Physical Sciences Division, the NASA Human Research Program, and Intel through a 2021 Frontier Development Laboratory Astronaut Health challenge. Cloud computing resources via Amazon Web Services were provided through a NASA SPARK award to Graham Mackintosh. HPE’s Spaceborne Computer is a collaboration between HPE and the International Space Station National Lab, with support from NASA. The study at OSD-435 was funded by the National Space Biomedical Research Institute [RE03701 through NCC 9–58 to MB] and the National Aeronautics and Space Administration [80NSSC17K0425 to MB].

## Software and data availability

The software and data used to run these experiments is freely available at https://github.com/nasa/AI4LS/tree/main/FLUID. The dataset used in running CRISP on Ground and the International Space Station are from the NASA Open Science Data Repository, OSD-435, https://doi.org/10.26030/dc1p-q547.

